# Distinct spatial domains of auxin perception control root growth in Arabidopsis

**DOI:** 10.64898/2026.11.18.767646

**Authors:** Shiv Mani Dubey, Julian Elijah Politsch, Martina Schmidtová, Eva Medvecká, Pavel Krupař, Matyáš Fendrych, Krzysztof Wabnik

**Author notes:** These authors contributed equally (Shiv Mani Dubey, Julian Elijah Politsch).

## Abstract

Primary root morphogenesis is coordinated by the integrated action of cell division, expansion, and differentiation. Although auxin is a central regulator of root development, how the spatial context of auxin perception shapes growth dynamics remains poorly understood. Here, we implement a synthetic auxin perception system that enables tissue-specific manipulation of auxin signaling output in defined spatial domains of the *Arabidopsis thaliana* root. Using this approach, we show that root growth depends primarily on the integration of local auxin perception in elongating cells of the outermost root tissue layers. Computational modeling predicts that mutually exclusive auxin sensitivity domains shift the position of the cell proliferation boundary, which was validated experimentally. We further found that elongating cells exhibit greater sensitivity to auxin input than meristematic cells, and that this differential responsiveness is critical for determining the overall root length. Together, our results identify mutually exclusive spatial domains of auxin sensitivity that coordinate root morphogenesis.

## Introduction

The precise regulation of plant cell division and elongation across tissues and organs is essential for proper plant growth, development, and pattern formation^1,2^. The *Arabidopsis thaliana* primary root provides an excellent model for studying these processes because of its highly organized cellular architecture and well-defined longitudinal spatiotemporal growth gradient^3,4,5^. Along the proximal–distal axis, the root is composed of a meristematic zone (MZ) of actively dividing cells, followed by an elongation zone (EZ) composed of elongating cells that have exited the cell cycle, and finally differentiating cells^4,5,6^. Radially, the root is organized into concentric tissue layers with specialized functions, including the lateral root cap (LRC), epidermis and cortex in the outer layers, and the endodermis, pericycle and vascular tissues in the inner cylinder^3,7^.

The phytohormone auxin is a central regulator of this spatiotemporal coordination during root morphogenesis. Several studies have demonstrated that local cellular responses to auxin steer developmental patterning in roots and other plant organs^8,9,10,11^. These responses emerge from the coordinated action of local auxin biosynthesis^12,13^, directional cell-to-cell auxin transport mediated by PIN and other transporters^7,14,15^, and auxin perception through the ubiquitously expressed TIR1/AFB family receptors and their downstream auxin signaling components, including auxin response factors (ARFs) and AUX/IAA transcriptional repressors^10,16,17^. Despite decades of intensive research in these areas, how the spatial organization of auxin biosynthesis, transport, perception, and signaling is integrated to produce robust and coordinated root growth remains incomplete.

Root growth depends on auxin responses operating across multiple timescales, encompassing both rapid and long-term regulatory mechanisms^6,8,18,19^. Elevated auxin levels inhibit cell expansion within minutes after hormone application^18,19,20^. This rapid response is essential for adaptive root bending during gravitropism, where asymmetric auxin accumulation differentially regulates cell elongation across the root^7,16,19^. Long-term auxin signaling is essential for maintaining meristem activity, as evidenced by the severe defects in dividing meristematic cells observed in auxin-resistant Aux/IAA mutants or following pharmacological inhibition of auxin perception^6,8,21,22^. Although root meristem size is defined by the extent of the proliferative domain^1,4,22^, much less attention has been paid to the cells located at the meristem– elongation zone interface that have largely exited mitosis but have not yet fully elongated. Recent work demonstrating that mechanical push from elongating tissues can reposition the meristem boundary further suggests that these transition-stage cells may actively contribute to meristem organization through non-cell-autonomous mechanisms^23^. However, how auxin perception in distinct tissues and developmental domains contributes to root morphogenesis remains unknown. To directly link local auxin responses with developmental outcomes, only a limited number of studies have manipulated auxin biosynthesis or auxin signaling in specific root tissues^7,8,24^. Tissue-specific expression of an auxin biosynthesis gene showed that auxin production in both inner and outer tissues influenced root growth, with the strongest effects arising from the epidermis and endodermis^8,24^. Ding et al.^8^ combined tissue-specific auxin biosynthesis with expression of the stabilized Aux/IAA17 (*axr3-1*) protein to either enhance or repress auxin responses in individual tissues, revealing that perturbations in both inner and outer tissues generally produce significant effects when compared to wildtype. However, tissue-specific auxin biosynthesis is inherently convoluted by auxin transport between neighboring cells, making it difficult to distinguish local signaling from auxin redistribution^8,24^. Moreover, stabilized Aux/IAA proteins artificially lock auxin signaling into constitutively repressed states and therefore do not preserve the native dynamics of auxin-responsive transcription^8,10^. Consequently, it remains unclear how individual tissues contribute to root morphogenesis when auxin perception is active while allowing the endogenous signaling machinery to operate normally.

Here, we combined an orthogonal auxin perception system with spatio-temporal computer modeling to dissect tissue-specific auxin responses at an unprecedented level of precision. While the ccvTIR1–cvxIAA system^25^ enabled selective activation of endogenous auxin signaling in defined tissues without perturbing auxin transport, the computational model^26^ allowed us to independently manipulate tissue auxin sensitivity by simulating auxin-induced growth mechanics while controlling for tissue-specific receptor expression. Using this approach, we identified a previously overlooked population of distal meristem cells at the meristem–elongation interface that controls the meristem boundary dynamics. We also demonstrate that outer tissues possess intrinsically greater auxin sensitivity than inner tissues. Together, our results reveal a hierarchical organization of auxin sensitivity that coordinates root development by establishing distinct spatial domains of cell elongation and proliferation.

## Results

### Native auxin responses alter meristem-associated marker expression

Understanding how cell division and elongation are spatially coordinated is important for understanding root development. These developmental processes are often inferred from morphological features such as cell size, cell shape, and division activity, but such features can be altered by auxin itself, which strongly affects cell elongation and the spatial extent of root growth^1,27^. Although the root-cap boundary provides an anatomical landmark for assessing the extent of the proliferative cell population, its precise identification requires careful visualization of root-tip anatomy^4,28^. Markers of developmental states therefore provide a useful complementary approach for defining root growth domains independently of cell geometry. We used pKNOLLE, an established marker of actively dividing cells^29^, together with pK890 (AT5G46890), a differentiation marker we developed that becomes active in elongating cells (see Methods). Their complementary expression provides a molecular readout of the transition from cell division toward differentiation reprogramming while cell elongation.

First we examined the response of the meristem-associated pKNOLLE marker (pKnolle::ccvTIR1-mVenus; pKNOLLE line) and the cell differentiation-associated pK890 marker (pK890::ccvTIR1-mVenus; pK890 line) lines to exogenous indole-3-acetic acid (IAA). As ccvTIR1 does not bind native IAA^25^, these lines are useful to understand the native IAA responses. We applied a low concentration of IAA (10 nM) for 24 h to provide a mild increase in auxin input. Although this concentration has been reported to inhibit root growth^18^, we asked whether it was sufficient to alter the spatial distribution of either marker along the longitudinal root axis. Neither pKNOLLE nor pK890 showed significant change in the position of their expression domains following IAA treatment (Fig. 1A, B). We further quantified meristem size using the root-cap-based criterion, in which the distance between the quiescent center and the root-cap boundary provides an independent spatial measure of the meristem domain. Consistent with the marker analysis, 10 nM IAA did not significantly alter this measure of meristem size (Fig. 1C).

**Figure 1.**
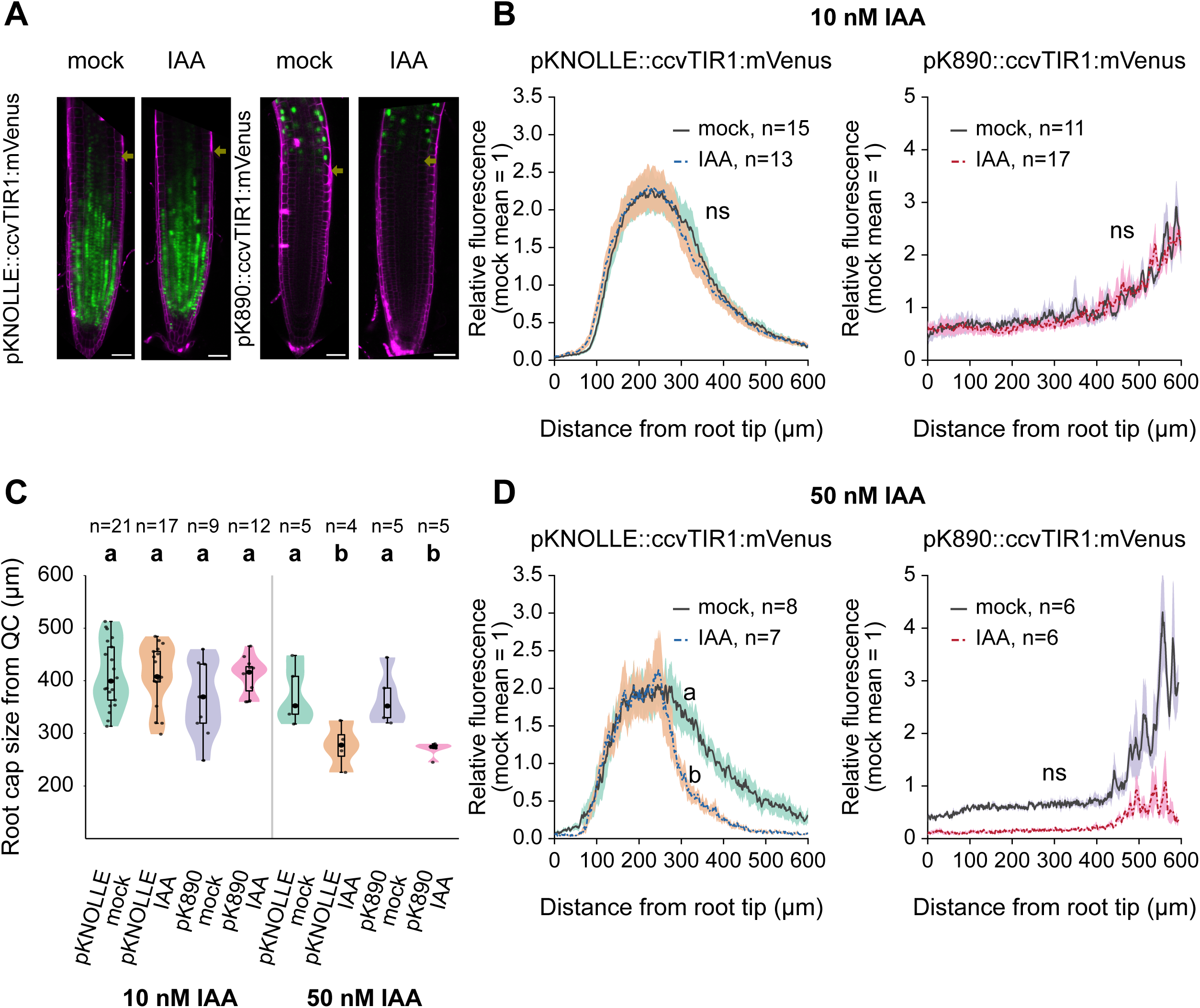
IAA treatment shifts the pKNOLLE division-zone marker toward the root tip in a dose-dependent manner, without significantly affecting pK890. **(A)** Representative confocal images of propidium iodide (PI)-stained roots (magenta) expressing pKNOLLE::ccvTIR1:mVenus or pK890::ccvTIR1:mVenus (green) under mock or 10 nM IAA treatment. Arrows indicate the root cap boundary. Scale bars, 50 µm. **(B)** Fluorescence intensity profiles of pKNOLLE::ccvTIR1:mVenus (left) and pK890::ccvTIR1:mVenus (right) along the longitudinal root axis, mock vs 10 nM IAA, normalized to the mock mean (see Methods). Mean ± SEM; n = number of roots, pooled from two independent experiments (pKNOLLE: 15 mock, 13 IAA; pK890: 11 mock, 17 IAA). Centroid positions were compared between treatments (Mann-Whitney U test); no significant shift (pKNOLLE: p = 0.49; pK890: p = 0.054; ns). **(C)** Root cap size, measured as the distance from the quiescent center (QC) to the root cap boundary (arrows in A), in pKNOLLE and pK890 lines under mock conditions and following treatment with 10 nM or 50 nM IAA. Violin plots show the data distribution; boxplots indicate the median and interquartile range; points represent individual roots (sample sizes indicated above each violin). Mock and IAA treatments were compared within each genotype using the Wilcoxon rank-sum test. Groups sharing the same letter are not significantly different. **(D)** As in (B), for 50 nM IAA from a single experiment (n = 8 mock, 7 IAA for pKNOLLE; 6 mock, 6 IAA for pK890). The pKNOLLE centroid shifted toward the root tip under IAA (65 µm, 95% CI 15-78 µm; p = 0.0037; letters a/b), while pK890 showed no significant shift (p = 0.70; ns).

We next asked whether a stronger auxin stimulus could alter the spatial distribution of these markers. Auxin levels have previously been shown to regulate meristem size and the balance between cell division and differentiation in the root, with the outcome depending on auxin concentration^21,22^. We therefore treated seedlings with 50 nM IAA for 24 h. Under these conditions, the pKNOLLE expression domain shortened toward the root tip, indicating that the spatial distribution of the meristem-associated marker is responsive to increased auxin input (Fig. 1D). In contrast, the pK890 expression profile did not show a significant positional shift following the same treatment (Fig. 1D). However, both the markers show reduction in the size of their root cap from QC following treatment with 50 nM IAA (Fig. 1C).

The different responses of the two markers may reflect the distinct developmental processes and temporal windows that they report. As shown below, the pKNOLLE and pK890 expression domains overlap at the meristem–elongation zone interface (Fig. 3C), suggesting that pK890 expression begins around the onset of cell elongation and differentiation. Because the treatment was maintained for 24 h, cells that initially entered the pK890 expression window may have progressed beyond this developmental stage by the time of imaging, potentially reducing the apparent pK890 signal. Thus, global exposure to 50 nM IAA may alter the timing or duration of the pK890 expression window through auxin-dependent transcriptional changes, without producing a detectable positional shift in its overall expression domain. Together, these findings establish that the cell division and differentiation markers can report auxin-dependent changes in the spatial organization of the root while also highlighting a limitation of global auxin applications: a whole-root auxin response cannot not reveal where auxin perception is primarily acting to produce the observed developmental outcome. Because of this limitation we decided to use an alternative strategy by controlling auxin output locally in domains defined by pKNOLLE and pK890 markers.

### High auxin sensitivity in elongating cells triggers root growth inhibition

To trigger auxin sensitivity boost in elongating and dividing spatial domains along the longitudinal axis of the root that facilitates auxin-dependent growth arrest, we implement a synthetic auxin perception system ccvTIR1-cvxIAA^18,25^ (Supplementary Fig. S1A). We expressed ccvTIR1 receptor responsive to synthetic cvxIAA broadly in the root using the native pTIR1 promoter (pTIR1::ccvTIR1-mScarlet; pTIR1 line) as global control or alternatively using pK890 and pKNOLLE lines(Fig. 2A). Seedling of pK890 genotype treated with cvxIAA for 4 hours caused inhibited root growth nearly as potent as pTIR1 line (Fig. 2B). However, this growth inhibition was marginal in the pKNOLLE line (Fig.2B). These data reveal that boosting auxin sensitivity specifically in different parts of the root has radically different impact on the root growth dynamics.

**Figure 2.**
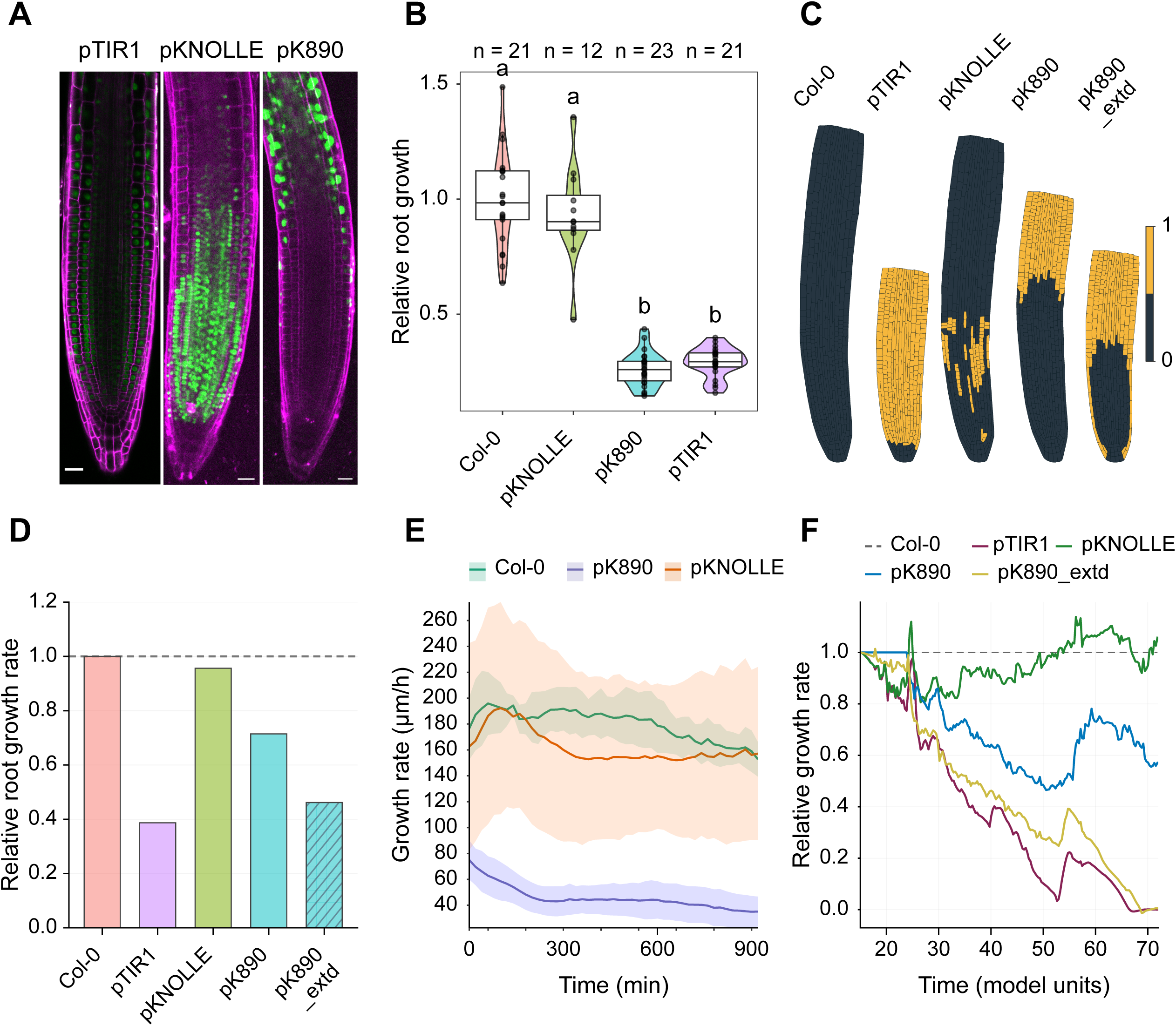
Auxin perception in the elongation zone is sufficient to mediate rapid root growth inhibition. **(A)** Representative confocal images of propidium iodide (PI)-stained roots expressing ccvTIR1-mVenus under the meristem-specific pKNOLLE or the elongation zone-specific pK890 promoter, or ccvTIR1-mScarlet under the native TIR1 promoter (pTIR1). The pTIR1 image was linearly unmixed to separate mScarlet from PI, which lowers the apparent signal; its intensity is therefore not comparable with the other lines (see Fig. 3A for a non-unmixed root). Green fluorescence indicates promoter activity and the spatial domain of ccvTIR1 expression. Scale bars, 25 µm. **(B)** Violin plots with embedded box plots showing relative root growth after 4 h treatment with 500 nM cvxIAA normalized to the corresponding mock-treated roots. Each dot represents an individual root. Center lines indicate medians, boxes represent interquartile ranges, and whiskers indicate the data range. Numbers above the plots indicate the number of roots analyzed for each genotype. Statistical significance was determined using non-parametric one-way ANOVA followed by multiple comparisons. Different letters denote statistically significant differences (P < 0.05). **(C)** Two-dimensional simulations of the spatial domains of the ccvTIR1 cell level response in Col-0, pTIR1, pKNOLLE, pK890, and pK890_extd. Yellow indicates cells affected by simulated ccvTIR1, and dark blue indicates cells outside the domain. The extended pK890 domain includes an additional region in the lateral root cap. **(D)** Quantification of mean root growth of the model relative to Col-0 from model time 15 to 72. Bars represent mean growth rate for that condition normalized to Col-0 growth rate over the same time interval. **(E)** Time-course analysis of root growth rates following transfer to medium containing 500 nM cvxIAA. Roots were continuously imaged during vertical growth on auxin-containing medium. Solid lines represent mean growth rates and shaded ribbons indicate ± SD. Colors correspond to individual genotypes. **(F)** Time-course analysis of the simulated root growth rates. Growth rates between consecutive time points measured as axial extent of root divided by elapsed model time, then normalized to the Col-0 growth rate over the same time interval. A value of 1 indicated the same growth rate as Col-0 simulation.

To further understand how observed root growth inhibition could be explained by local differences in auxin sensitivity across the longitudinal root axis, we employed a spatio-temporal computational model representing realistic 2D root growth dynamics, built on the foundational models^5,23,26,30^. Our model implements different spatial domains of boosted auxin perception where the synthetic auxin sensing is activated following the experimentally observed ccvTIR1 expression (Fig. 2A,C, Supplementary Movie S1A). We model cell wall stiffness as a function of cellular auxin input throughout the model, and defined synthetic domains increase cellular stiffness above baseline. (see Methods,^26^). Our simulations predicted that boosting auxin perception in elongating regions of the root in the simulation of pK890 line led to moderate overall inhibition of total root growth (Fig. 2D), in part due to meristematic overgrowth compared to modeled Col-0; however, despite this moderate reduction in overall root growth, the simulated elongation-zone size was reduced to a degree comparable to that observed in the pTIR1 model (Fig. 4E). To investigate this discrepancy between model prediction and observations, we monitored root growth dynamics by long-term live imaging using a specialized imaging chamber mounted on a vertical microscope (see Methods;^20^). Over the 900 min imaging period, the pK890 line exhibited sustained inhibition of root growth, whereas the pKNOLLE line showed an initial reduction in growth rate followed by a recovery to its normal growth rate (Fig. 2E). Noticeably, the ccvTIR1-mVenus fluorescence in the pK890 line was detected not only in the elongation zone but also dynamically extended toward the proximal meristem, particularly in the lateral root cap (LRC) (Supplementary Fig. S2A; Supplementary Movie S2). Based on these intriguing observations, we modified the ccvTIR1 expression domain in our pK890 model to include LRC cells which we call the pK890_extd model phenotype (Fig. 2C). Indeed, the adapted model predicted a substantially stronger inhibition of root growth (Fig. 2D; Supplementary Movie S1B) together with a sustained inability to recover normal growth dynamics following auxin input (Fig. 2F). In the case of meristematic ccvTIR1 activation in the simulated pKNOLLE line, we observed a mild root growth inhibition (Fig. 2D, F; Supplementary Movie S1B).

Together, these simulations and data indicate that pace of root growth depends on the state of auxin sensing within a spatial domain predominantly occupied by elongating cells. In contrast, auxin signaling in actively dividing meristematic cells has weak transient impact on root elongation dynamics, highlighting a functional separation between these two cellular domains.

### Auxin signaling in elongating and dividing cells jointly contribute to a dynamic meristem boundary positioning

Our findings indicate that auxin perception and signaling in elongating cells modulates root growth dynamics, which agrees with the high intrinsic strain rates of this developmental domain^31^. However, root meristem activity depends on a balance between cell division and cell expansion. Although auxin signaling is a key regulator of meristem function, and elevated auxin levels can markedly reduce meristem size^6,22,41^, it remains unclear whether auxin controls meristem size primarily through its direct signaling activity within the meristematic domain or also indirectly through its effects on cell elongation in adjacent developmental domains. To address this question, we quantified meristem size following a 20h treatment with cvxIAA in the pKNOLLE, pK890, and pTIR1 lines. Activation of synthetic auxin perception throughout the root in the pTIR1 line caused a pronounced reduction in meristem size (Fig. 3A,B). Notably, activation of auxin signaling in elongating cells of the pK890 line also substantially reduced meristem size, to an extent comparable to that observed in the meristem-specific pKNOLLE line (Fig. 3A,B).

**Figure 3.**
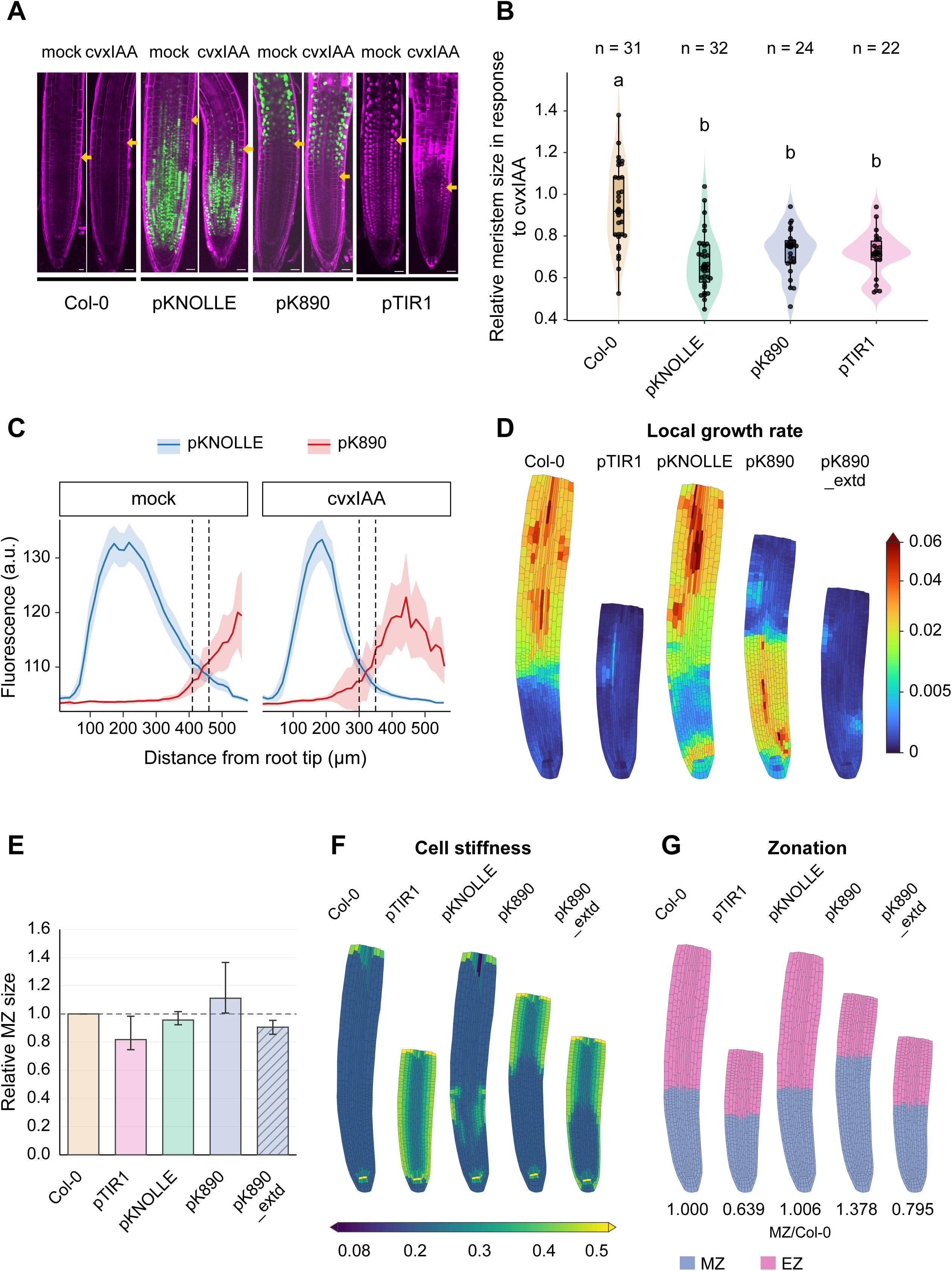
Auxin signalling in the intersection domain of MZ and EZ decides the meristem boundary and is recapitulated by computational modeling. **(A)** Representative confocal images of propidium iodide-stained roots expressing pKNOLLE:ccvTIR1:mVenus, pK890::ccvTIR1:mVenus, and pTIR1::ccvTIR1:mScarlet treated with mock or 500 nM cvxIAA for 20 h. Yellow arrowheads indicate the position of the meristem boundary. Scale bars, 50 μm. **(B)** Violin plots with embedded box plots showing relative meristem size normalized to the corresponding mock-treated roots of each genotype. Individual dots represent single roots. Center lines indicate medians, boxes represent interquartile ranges, and whiskers indicate the data range. Numbers above the plots indicate the number of roots analyzed for each genotype. Statistical significance was determined using non-parametric one-way ANOVA followed by multiple comparisons. Different letters indicate statistically significant differences (P < 0.05). **(C)** Fluorescence intensity profiles of meristematic and elongation zone markers measured along the longitudinal axis from the root tip in mock- and cvxIAA-treated roots. Solid lines represent mean fluorescence intensity and shaded ribbons indicate ± SE. Dotted lines mark the intersection between both marker domains, corresponding to approximately 420 μm in mock-treated roots and 315 μm in cvxIAA-treated roots, indicating an apical shift of the meristem–elongation zone transition. **(D)** Cell level map showing growth rate in selected phenotypes in simulated roots. Colors indicate the rate of increase in cell area. Dark blue indicates little to no growth and red indicates rapid cell growth. Values are normalized by square root to show useful ranges. Units are relative to model and arbitrary. Triangles in **D** and **F** mark values beyond the color limits. **(E)** Relative meristem length of simulated roots measured as the span of meristematic epidermal cells along the roots axis, and normalized to Col-0 at each time point. Bars show the mean ratio from meristem appearance in model simulation (time 30) to end of simulation. Whiskers span the highest and lowest ratio for each phenotype with Col-0 during that period. **(F)** Spatial distribution of cell stiffness along the root axis and across tissue layers in selected root simulations. Cells are colored by stiffness; higher values indicate greater resistance to deformation and growth pressure. **(G)** Simulated model root zonation. Blue indicates meristematic zone (MZ) and pink indicates elongation zone (EZ). Values below each root phenotype indicate relative distance from quiescent center (QC) to the lateral root cap (LRC) bounty in simulated phenotype compared to Col-0.

To further investigate how auxin signaling in dividing and elongating cells influences the position of the meristem boundary, we quantified the spatial distribution of ccvTIR1–mVenus fluorescence along the primary root axis in the pKNOLLE and pK890 lines. Under mock conditions, the fluorescence profiles intersected approximately 400–450 μm from the root tip, corresponding to the transition between the meristematic zone and the advancing front of elongating cells (Fig. 3C). Following cvxIAA treatment, this intersection shifted closer to the root tip (approximately 300–350 μm), indicating an apical displacement of the meristem–elongation boundary (Fig. 3C).

Live imaging further revealed that mVenus fluorescence in pKNOLLE line progressively disappeared toward the distal meristem, consistent with a decline in mitotic activity near the end of the meristem boundary (Supplementary Movie S2). In contrast, the mVenus fluorescence in pK890 lines is apparently scattered in LRC within this region. (Supplementary Movies S2 and S3). Together, these observations suggest that the position of the meristem boundary is determined not only by the extent of cell proliferation but might also be due to auxin signalling in elongating cells and LRC cells. To further explore whether auxin signalling in elongating cells is sufficient to determine the meristem boundary or mechanical perturbations imposed by LRC on inner tissues in meristem region is crucial for the experimentally observed changes in meristem size, we simulated the pK890 model phenotype to test whether inhibition of cell elongation would lead to meristem shrinkage, as observed experimentally, or instead promote meristem expansion, as recently proposed ^23^. In the pK890 line scenario the model predicted a modest enlargement of the meristem relative to the control (Fig. 3D and E, Supplementary Movie S1D), consistent with previous reports ^23^. However, this result could not explain the experimentally observed reduction in meristem size in the pK890 line (Fig. 3B). We therefore used the extended pK890 model phenotype to include experimentally observed LRC cells (pK890_extd scenario; Fig. 2C, Supp. Movie 1A). Again, under these conditions, the model predicted meristem shrinkage, closely recapitulating the phenotype observed experimentally in the pK890 line (Fig. 3D-G; and Supplementary Movies S1 and S3). While the pK890_extd scenario resulted in sustained model root growth inhibition across the root over time (Fig. 2F), model root growth inhibition in the pKNOLLE scenario gradually diminished owing to reduced meristematic activity and progressive contraction of the meristem region (Fig. 3D-G).

Collectively, these findings reveal auxin signaling in the elongation zone does not appear to determine the position of the meristem boundary directly. Instead, meristem organization appears to be governed by auxin signaling within the meristematic domain, with mechanical constraints imposed by LRC cells further contributing to the positioning of this boundary. Since pK890_extd model phenotype does not show reduction in meristem size up to the comparable level to pTIR1 model phenotype, suggesting apart from mechanics, a possible crosstalk with other phytohormones downstream of orthogonal auxin perception and signalling, may also contribute to meristem organisation.

### Boosting auxin sensitivity in elongating cells exclusively controls the span of elongation region

We therefore investigated how mutually exclusive domains of auxin perception (in elongating versus dividing cells) influence the spatial organization of the root elongation domain. We speculated that enhanced auxin perception in dividing cells could potentially accelerate the transition toward differentiation through auxin-mediated transcriptional reprogramming^6,9^, thereby indirectly affecting the size of the elongation region.

To test this, we quantified the elongation zone length after 20 h of cvxIAA treatment (Fig. 4A,B). The size of the elongation zone was determined as the distance between the intersection of the meristematic and elongation-zone marker domains and the onset of differentiation. In wild-type roots, this transition occurred approximately 420 μm from the root tip, whereas in cvxIAA-treated roots the transition shifted closer to the apex, to approximately 315 μm (Fig. 3C). Consequently, the pKNOLLE line exhibited less reduction in elongation zone size compared to the reduction observed in the pK890 and pTIR1 lines (Fig. 4B). These observations suggest that auxin signaling in the meristem alone has a limited influence on elongation zone size and cannot account for the strong inhibition observed when auxin signaling was boosted in elongating cells.

**Figure 4.**
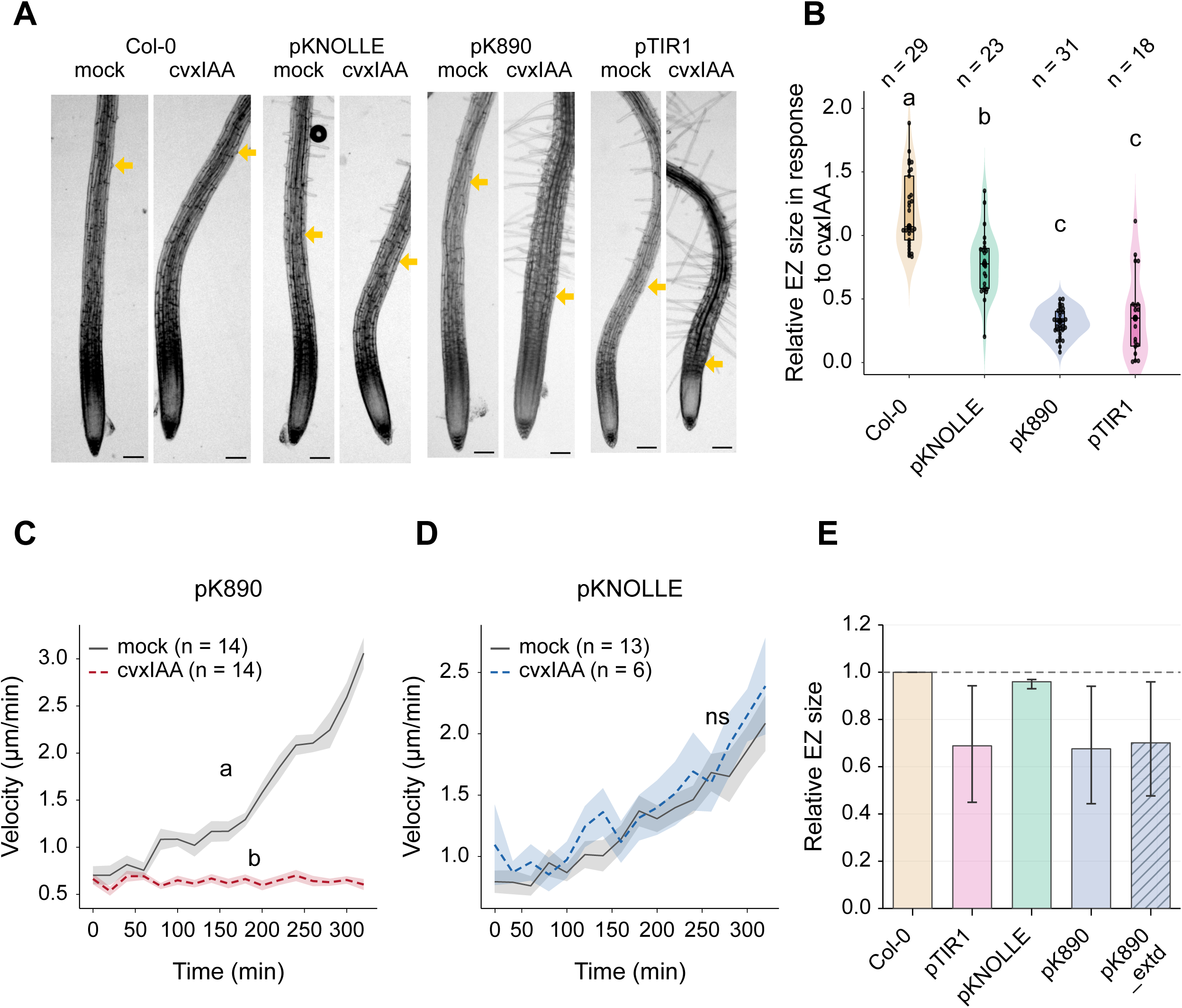
Cell type-specific auxin signaling regulates elongation zone size and cell displacement in the Arabidopsis root. **(A)** Representative root images showing the effect of 20 h treatment with 500 nM cvxIAA on elongation zone (EZ) size in Col-0, pKNOLLE::ccvTIR1, pK890::ccvTIR1, and pTIR1::ccvTIR1 roots. Yellow arrows indicate the position of the first root hair bulge, marking the transition between the elongation and differentiation zones. Scale bar = 100 µm. **(B)** Quantification of the relative elongation zone response to cvxIAA treatment. Elongation zone length was determined as the distance between the pKNOLLE and pK890 expression domain boundaries. In Col-0 roots, the elongation zone measured approximately 420 µm under mock conditions and 315 µm following cvxIAA treatment. Values were normalized to the corresponding mock-treated control of each genotype. Numbers above violin plots indicate the number of roots analyzed. Statistical significance was assessed using non-parametric ANOVA. **(C)** Nuclear velocity during epidermal cell elongation in the *pK890::ccvTIR1:mVenus* line. Nuclei were manually tracked in Fiji/ImageJ from time-lapse movies of root growth under mock or 500 nM cvxIAA treatment; nuclear displacement between consecutive 20 min time points was used to calculate velocity (displacement/time interval). Lines and shaded bands show mean ± SEM velocity at each time point over the 0–320 min window, during which tracking data were available for all nuclei in both conditions (mock: n = 14 nuclei from 3 roots; cvxIAA: n = 14 nuclei from 3 roots). Per-nucleus mean velocity over this window was compared between mock and cvxIAA by Mann–Whitney U test. **(D)** As in (C), for the *pKNOLLE::ccvTIR1:mVenus* line (mock: n = 13 nuclei from 3 roots; cvxIAA: n = 6 nuclei from 2 roots). Unlike pK890, pKNOLLE cvxIAA-treated cells showed no significant change in elongation velocity relative to mock in Mann–Whitney U test. **(E)** Relative elongation zone (EZ) length in simulated roots. Length was estimated by the span of EZ epidermal cells along the roots axis and normalized to Col-0 at each time point. Bars show the mean ratio over model time 30-72. Error bars span the minimum and maximum ratio during that period.

The moderate reduction in elongation zone size in pKNOLLE line indicates several possibilities. Cells exiting the meristem could (i) elongate at the same rate but initiate differentiation earlier, or (ii) elongate more slowly and initiate differentiation earlier. To distinguish between these two possibilities, we followed individual cells over time and quantified their velocity during their elongation (Fig. 4C). In the pK890 line, activation of auxin signaling caused a pronounced reduction in cell elongation velocity, demonstrating that local auxin signaling boost in elongating cells directly limits cell expansion (Fig. 4C). In contrast, cells in the pKNOLLE line showed no significant change in velocity relative to mock treatment (Fig. 4D). Notably, the elongation velocity of cvxIAA-treated pKNOLLE cells was statistically indistinguishable from that of mock treated pK890 cells (Supplementary Fig. S2B), indicating that these cells elongate at a rate comparable to unperturbed elongation. This is consistent with the possibility (i): pKNOLLE-triggered cells elongate at a normal rate but exit the elongation zone and differentiate sooner, rather than elongating more slowly over a prolonged window (possibility ii). Together with the modest reduction in elongation zone size (Fig. 4B), this indicates that the weak effect of meristem-specific signaling on root growth reflects an earlier transition out of elongation, not impaired elongation kinetics.

To further assess whether meristem-derived auxin responses could indirectly influence elongation, we simulated domain-specific auxin inputs with our computational model. Consistent with the experimental observations, the model predicted only a modest reduction in EZ when auxin signaling was boosted in dividing cells, whereas activation in elongating cells or throughout the root caused a much stronger reduction in cell movement and elongation rates (Fig. 4E, Fig. 3D-F, Supplementary Movie S1D). These simulations closely matched the experimentally measured changes in elongation zone size (Fig. 4B), further supporting the biological plausibility of our proposed model.

Together, these findings indicate that auxin signaling in the meristem and elongation zone controls distinct developmental outputs. Auxin responses in dividing cells primarily regulate meristem organization and cell production, whereas local auxin perception within elongating cells predominantly determines cell expansion dynamics and therefore defines the extent of the elongation zone. Thus, the size of the elongation zone is controlled locally by auxin-mediated regulation of cell elongation rather than by meristem-derived signals.

### Outer root tissues dominate auxin-mediated growth inhibition

Building on our finding that rapid root growth inhibition is predominantly mediated by auxin signaling in the elongation zone, we next asked whether specific tissues along the radial axis differ in their contribution to this response. Using the tissue-specific ccvTIR1 lines, we quantified root growth following a 4 h cvxIAA treatment. Activation of auxin signaling in outer tissues–particularly elongating trichoblasts (pCOBL9 line), cortex cells (pPEP line), and the broader PIN2 domain (pPIN2 line) encompassing the epidermis, cortex, and lateral root cap–resulted in significant growth inhibition compared with wild type (Fig. 5A,B). Among these, activation in the PIN2 domain produced the strongest inhibitory effect, although it remained weaker than that observed in the ubiquitous pTIR1 line, suggesting that maximal growth inhibition requires the combined contribution of multiple tissues.

**Figure 5.**
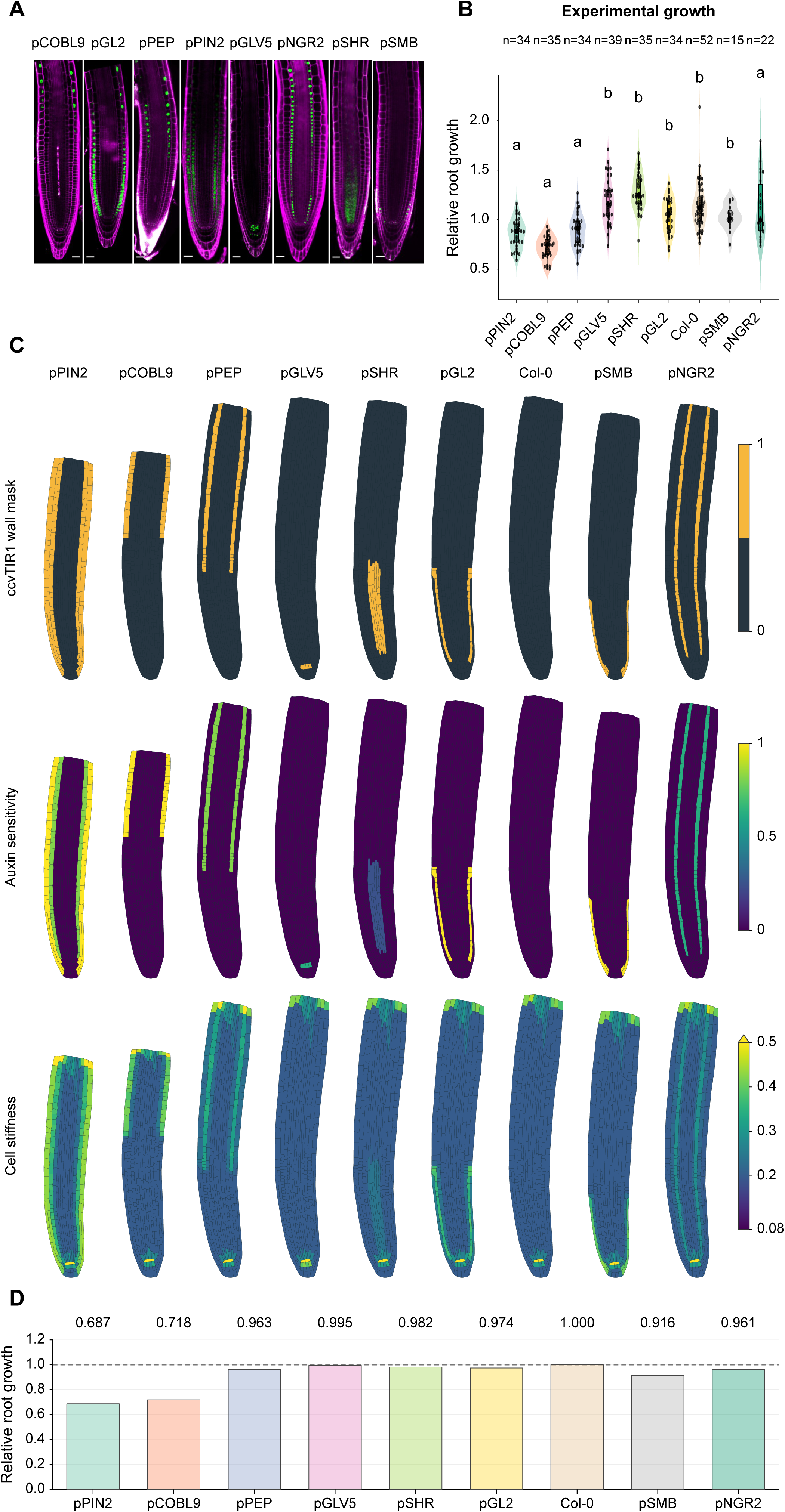
Root growth inhibition is majorly controlled by highly sensitive outer root tissues. **(A)** Representative propidium iodide-stained roots showing tissue-specific expression domains of ccvTIR1 driven by pCOBL9 (trichoblasts), pGL2 (atrichoblasts), pPEP (cortex), pPIN2 (epidermis, cortex, and lateral root cap), pGLV5 (columella), pNGR2 (endodermis), pSHR (stele), and pSMB (lateral root cap). Scale bar = 25 µm. **(B)** Quantification of root growth following treatment with 500 nM cvxIAA for 4 h. Root growth values were normalized to the corresponding mock-treated control for each genotype. Numbers indicate the number of roots analyzed per genotype. Different letters denote statistically significant differences as determined by non-parametric ANOVA. **(C)** Model root maps of the ccvTIR1 response domain (top), auxin sensitivity (middle), and cell stiffness (bottom) for each simulated condition. In the top row, yellow indicates cells within the response domain, followed by assigned auxin sensitivity in the defined domain in the middle row, followed by physiological outcome in terms of cellular stiffness in the bottom row. **(D)** Relative root growth rate in the simulations from model time of induction of ccvTIR (15) to end of simulation (72). The mean growth rate was calculated as the change in root length divided by elapsed time. Bars show this rate for each phenotype relative to the growth rate of Col-0 over the same time interval.

In contrast, activation of auxin signaling in inner tissues, including the stele (pSHR line), columella and epidermal initials (pGLV5 line), and the lateral root cap/inner tissue domain (pNGR2 line), did not inhibit growth and in some cases showed a modest trend toward growth promotion (Fig. 5B). As these promoters differ not only in their radial expression domains but also in their longitudinal expression patterns along the root axis, the observed phenotypes likely reflect the combined influence of both spatial dimensions. Nevertheless, the results are consistent with stronger growth-inhibitory responses arising from outer tissues compared with the inner tissue domains examined. Consistent with the longitudinal analyses, auxin activation in atrichoblast and meristem-associated domains had no significant effect on short-term root growth, further supporting the conclusion that rapid auxin-dependent growth inhibition is mediated primarily by outer elongating tissues. Collectively, these findings demonstrate that auxin-mediated root growth inhibition is not uniformly distributed across the root but is instead dominated by the epidermis– cortex axis, with the greatest inhibition achieved through coordinated signaling across multiple outer tissues.

To determine whether this hierarchy arises solely from organ-level mechanics or because outer tissues possess intrinsically greater sensitivity to auxin-mediated growth regulation, we performed simulations in which tissue-specific ccvTIR1 expression domains were introduced radially and tissue sensitivity to auxin-like cell wall stiffness effects were tested independently. When all tissues were assigned equal auxin sensitivity, the model failed to reproduce the experimentally observed hierarchy and instead predicted substantial contributions from both outer and inner tissues (Supplementary Movies S4 and S5). Similarly, increasing the sensitivity of inner tissues generated growth inhibition patterns that were inconsistent with the experimental data (Supplementary Movie S5). In contrast, assigning greater auxin sensitivity to outer tissues successfully recapitulated the experimental hierarchy, correctly predicting stronger growth inhibition following PIN2 domain-, cortex-, and trichoblast-specific perturbations (Fig. 5C and D, Supplementary Movie 4A-D).

Furthermore, we asked whether a similar hierarchy exists during long-term root morphogenesis by experimentally quantifying the relative contributions of individual tissues to meristem and elongation domains. In the meristem, pPIN2 roots displayed a significant reduction in meristem size compared with Col-0, whereas GL2 roots exhibited a tendency toward meristem reduction that did not reach statistical significance (Supplementary Fig. S3A). Notably, these trends were closely reproduced by the computational model (Supplementary Fig. S3C; S5F). Analysis of elongation zone size revealed a similar tissue hierarchy to that observed during the short-term root growth response (Supplementary Fig. S3B). Specifically, auxin signaling in the outer tissues exerted the strongest effect on elongation zone development, with pCOBL9, pPIN2, and pPEP lines displaying the most pronounced reductions in elongation zone size (Supplementary Fig. S3B). Importantly, this radial pattern was closely recapitulated by the computational model, which similarly predicted a dominant contribution of outer tissues to elongation zone regulation (Supplementary Fig. S3D; S5D). Together, these findings indicate that the hierarchical contribution of root tissues to auxin-mediated morphogenesis extends beyond rapid growth inhibition and governs the long-term establishment of root zonation.

Furthermore, we tested the three sensitivity scenarios for pK890, pKNOLLE, pTIR1 as well as pK890_extd lines in the three tissue sensitivity scenarios. The model most closely predicted the experimentally observed data in the simulation with high outer sensitivity regarding root growth, EZ size, and MZ size. (Supplementary S5C,E and G). These simulations suggest that the dominant role of outer tissues cannot be explained solely by their position within the organ or by tissue mechanics. Rather, they indicate that outer tissues possess intrinsically higher responsiveness to auxin-mediated growth regulation. Interestingly, the mild growth-promoting tendency observed in stele- and columella-specific lines may reflect tissue-specific functions of auxin that are distinct from growth inhibition, or alternatively indirect effects on endogenous auxin distribution and homeostasis. Although the underlying mechanism is unclear, these observations further support the notion that outer tissues are both the most auxin-sensitive and the most effective mediators of rapid auxin-dependent root growth inhibition.

## Discussion

Auxin is a master regulator of plant growth and development and is involved in multiple phytohormone pathways through extensive crosstalk at the levels of biosynthesis, transport, and signaling^32^. Although spatial auxin gradients are widely considered an important determining factor in developmental patterning, it remains unclear whether all tissues contribute equally to auxin-mediated morphogenesis. Previous studies addressing tissue-specific auxin function relied either on local auxin biosynthesis^8,24^ or on constitutive modulation of auxin signaling through dominant Aux/IAA proteins^8,33^. However, auxin transport complicates the interpretation of local biosynthesis experiments, while dominant Aux/IAA approaches do not isolate auxin perception itself and often impose binary effects on transcriptional output. By employing the engineered orthogonal ccvTIR1-cvxIAA system, we boosted auxin signaling in defined tissues independently of endogenous auxin transport, providing a framework to directly compare the relative contributions of individual tissues to root morphogenesis. An additional challenge in comparing tissue specific lines is that different promoters can drive different levels of ccvTIR1 expression, making it difficult to distinguish differences in tissue contribution from differences in receptor abundance. By combining our experimental approach with computational simulations in which auxin input was controlled across tissues, we could examine how tissue identity and responsiveness contribute to the observed growth responses.

Our results demonstrate that auxin-mediated root morphogenesis is organized hierarchically rather than uniformly across tissues. Although activation of auxin signaling in several tissues contributed to growth regulation, outer tissues, particularly the epidermis and cortex, exerted the strongest effects on both short-term root growth inhibition and long-term developmental effect. Importantly, the ubiquitous pTIR1 line consistently produced the most severe phenotypes, indicating that root morphogenesis results from integrated auxin response from several tissues rather than from a single dominant tissue.

The dominant contribution of outer tissues may result from several non-mutually exclusive mechanisms. Because outer cell layers directly determine organ surface expansion, they may mechanically constrain inner tissues during growth^34^. Additionally, tissue-specific compositions of ARFs and Aux/IAA proteins may generate differential auxin responsiveness across tissues^9,17^. Similar mechanical coordination has been described for epidermal brassinosteroid signaling^35,36^, suggesting that auxin signaling may also exert organ-wide effects through mechanically coupled outer tissues.

Importantly, our computational model allowed us to distinguish between tissue mechanics and intrinsic tissue responsiveness. Although differences in promoter strength here cannot be completely excluded experimentally, simulations assuming equal tissue sensitivity failed to reproduce the observed hierarchy. Instead, the model performed best when outer tissues were assigned greater auxin sensitivity than inner tissues. These findings suggest that mechanics alone are insufficient to explain tissue hierarchy and support the existence of differential auxin responsiveness among tissues.

Our results reveal that auxin outputs controlling cell production and cell elongation are spatially distinct yet interconnected. Auxin signaling restricted to dividing cells strongly influences meristem size, while signaling in elongating cells predominantly controls cell expansion and root growth. Notably, perturbations in one domain affect the organization of the other, demonstrating non-cell-autonomous communication between the meristematic and elongating cells. Beyond mechanical interactions, these responses may also involve crosstalk between auxin and other phytohormones, potentially leading to transcriptional reprogramming and changes in cell differentiation. Thus, root organisation does not appear to arise from independent spatial developmental modules, but rather from coordinated auxin responses across adjacent domains. An interesting finding is that meristem boundary positioning depends primarily on auxin signalling within the meristematic domain, whereas activation of auxin signalling specifically in the elongation zone does not directly alter meristem size. This suggests that the positioning of the meristem boundary is more closely associated with local auxin responses in the dividing region than with auxin signaling in the adjacent elongation zone.

Although the computational model captured several aspects of the experimental phenotypes, it did not fully reproduce some observed responses, including the increase in meristem size in the pCOBL9 condition. This discrepancy suggests that additional regulatory components may contribute to the observed phenotypes. In particular, the interaction between hormonal signalling and tissue mechanics may be more complex than currently represented in the model. Crosstalk with other phytohormones could provide an additional layer of regulation, potentially modifying mechanical properties and influencing the relationship between cell proliferation and elongation.

We show that meristem specific signaling does not have effect on the cell elongation rate after cells stop dividing and leaves the zone for elongation, interestingly, they elongate at the same rate, but their elongation zone size is small. A possible explanation is that they might have accelerated the differentiation reprogramming while being in the division phase and being exposed to extra auxin signalling by ccvTIR1-cvxIAA. In this scenario, cells would complete the differentiation program sooner leaving the elongation zone size small. However, experimental evidence is needed to support this explanation.

Together, our findings postulate a model in which root morphogenesis emerges from coordinated auxin responses distributed across spatially distinct domains. Rather than being governed by a single tissue or developmental zone, root growth and zonation result from unequal yet integrated auxin responses, with outer tissues and elongation associated cell expansion serving as dominant regulators. Future single-cell or single-nuclear transcriptomic analyses, along with combinations of auxin and mechanical reporters, will help define the molecular basis underlying tissue-specific auxin sensitivity and non-cell-autonomous coordination during root development.

## Materials and methods

### Plant materials and growth conditions

All *Arabidopsis thaliana* tissue-specific transgenic lines in this study were of Col-0 background. We used the homozygous stage of all ccvTIR1 lines except NGR2:ccvTIR1-mVenus and SMB:ccvTIR1-mVenus. Seeds were sterilized with bleach (3% NaClO) for 2 h, and stratified at 4 °C in the refrigerator for 2 days. Plants were grown vertically on ½ Murashige-Skoog (MS) medium (Duchefa) containing 1.0% agar (Duchefa), 1% (w/v) sucrose, adjusted to pH 5.8 with KOH, and maintained at 22 ^0^C in a long-day growth room (16L/8D) at 60% humidity and a light intensity of 100 mmol photons m -2 s -1.

### Molecular cloning

The TIR1 binding pocket was modified by the substitution mutations F79 > G to create ccvTIR1 ^25^. The following promoters were used for tissue-specific targeting of ccvTIR1-pTIR1 (native), pPIN2 (lateral root cap, epidermis, cortex), pCOBL9 (trichoblast epidermis), pGL2 (atrichoblast epidermis), pPEP (cortex), pNGR2 (endodermis), pSHR (stele), pGLV5 (columella), pKNOLLE (dividing cells in the meristematic region of the root, pK890 for differentiating cells). Promoter pK890 of gene AT5G46890 was selected based on predicted specific expression in differentiating cells^37^. ccvTIR1 and all promoters were first domesticated into the pUPD2 vector. Then, the tissue-specific ccvTIR1 transcriptional units were designed in pDGB1alpha1 by combining the promoters and a C-terminal mScarlet or mVenus tag. These transcriptional units were then combined with a Basta resistance cassette into the pDGB3omega1 binary vector. Cloning steps were performed using the GoldenBraid method^38^). (https://gbcloning.upv.es/). The binary constructs were transformed into *Agrobacterium tumefaciens* strain GV3101 by electroporation and subsequently transformed into Col-0 plants by floral dipping.

### Measurement of meristem and elongation zones

Meristem and elongation zone lengths were quantified from vertical spinning disk microscopy images using Fiji/ImageJ^39^. For pKNOLLE and pK890 marker lines, fluorescence intensity profiles were obtained along the longitudinal root axis using the Segmented Line tool with a line width spanning the entire root diameter. Fluorescence intensity was measured from the quiescent center (QC) toward the differentiation zone, and the meristem–elongation zone boundary was defined as the intersection point between the meristematic and elongation zone marker expression profiles. For lines lacking exclusive meristem or elongation zone markers, meristem size was estimated as the distance from the QC to the distal end of the lateral root cap. Elongation zone size was determined by measuring the distance between the QC and the position of the first visible root hair bulge and subtracting the average meristem length obtained from the corresponding mock and cvxIAA treated roots. Measurements were performed using Fiji/ImageJ, and all values are presented relative to the respective mock controls. To compare the effect of IAA at 10 nM and 50 nM on the pKNOLLE and pK890 lines, fluorescence was measured along the root axis from 0 to 600 µm from the root tip, in mock- and IAA-treated roots at each concentration. Because the two concentrations were imaged in separate experiments, values were normalized within each experiment (each marker line at each concentration). First, the mock roots were averaged at every distance from the root tip to obtain a mean mock profile. The mean of this profile over 0-600 µm gave a single normalization value. All mock and IAA roots from the same experiment were then divided by this value, so that the mean mock fluorescence equals 1.

### Root growth analysis

To measure the short-term (4h) effect of cvxIAA on root growth, vertically grown, five-day-old seedlings were imaged every 30 min after transplanting onto medium containing 0 nM or 500 nM cvxIAA using a vertically positioned flatbed scanner at 1200 dpi. Root growth was measured by tracking root tips with the segmented line tool in Fiji^19^. To measure the long-term (5 days) effect of cvxIAA on root growth, seeds were sown on medium containing 0 nM or 500 nM cvxIAA and stratified in the refrigerator for 2 days and then grown vertically for five days. Response to cvxIAA was quantified by measuring primary root length. The entire region below the hypocotyl was included in the root length analysis. Root growth response to cvxIAA was normalized for both long-term and short-term treatment by dividing the root length (for long-term treatment) or root tip growth (for short-term treatment) by the average root length or average root tip growth under control conditions. The response value 1 means that the treatment did not affect root growth.

### Live imaging of root

Long term live imaging of roots was performed using the PlantScope Xero system with a Yokogawa CSU-W1 spinning disk equipped with Zeiss Plan-Apochromat 20×/0.8 objective. Plants were placed into a custom-made chambered cover slide under agar slab (1/2 MS salts, 1% sucrose, 0.8% Plant Agar, Duchefa) with propidium iodide (1ug/ml medium). Roots were imaged every 15 minutes, and the root tip was tracked using the SlideBook software (3i Intelligent Imaging Innovations).

### Nuclear tracking

Nuclear dynamics were analyzed using Fiji/ImageJ^39^. Time-lapse image sequences were first corrected for sample drift using the StackReg plugin. Epidermal cells entering the elongation zone were identified by the nuclear ccvTIR1 signal (pK890 line), and 3–4 nuclei per root were manually tracked using the manual tracking plugin. In total, three roots were analyzed for each genotype and treatment condition. Images were acquired every 15 min and nuclei were followed for a total duration of 300 min. Nuclear positions obtained from the manual tracking plugin were used to calculate cell displacement and velocity over time. Displacement and velocity were determined at each 20 min interval using the coordinates generated by the Manual Tracking plugin, allowing changes in nuclear movement dynamics to be monitored throughout the 300 min imaging period. Displacement and velocity values represent the distance traveled by each nucleus over time.

### Statistical analysis and graphics

Root data was analysed using Fiji software. Statistical analyses were conducted using R software (R v4.0.2 and RStudio v1.4.1106). Plots were generated using ggplot in R software. Images were compiled using Inkscape (v1.0). The study included 2-4 biological replicates, all showing consistent results. The Sol 6.1 model was used to optimize Python3 code involved in figure alignment and assembly. Unity was used for root model streaming and imaging (Unity Version 6000.0.23f).

### Spatio-temporal root growth model

Root growth simulations were performed in the MorphoDynamX framework^40^ with image streaming to the Unity engine and are based on previously published 2D mesh modules^23,26^. The root was represented as a two-dimensional cellular complex in which each cell carries tissue identity, developmental zone state, auxin-related chemical variables and mechanical properties. Simulations coupled auxin transport, turgor-driven wall mechanics, mechanical growth, cell division, remeshing and developmental zone transitions. Mechanical relaxation was solved with the model’s Position-Based Dynamics solver, while chemical updates were integrated with an explicit Euler scheme. Because the baseline framework has been described previously^26^, we describe here only the additions used to simulate cvxIAA/ccvTIR-dependent tissue sensitivity.

Promoter-like response domains were encoded using binary wall-response and growth-response masks (Supplementary Fig. S4). Receptor expression was uniform within each active mask and zero outside it; a separate radial weight represented sensitivity. Domains included broad pTIR1 expression and tissue patterns corresponding to pKNOLLE, pSMB, pGL2, pCOBL9, pPEP, pNGR2, pSHR, pGLV5, pPIN2, pK890 and pK890_extd as well as related controls. For each cell *i*, the response mask was defined by:

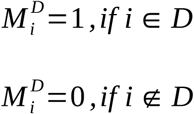

Here D is the selected ccvTIR expression domain. cvxIAA perception was activated only after a treatment delay, following experimental conditions, implemented in the parameter sweeps as cvxIAA delay. To avoid mechanical discontinuities, the response was introduced using a linear ramp over 10 simulation steps:

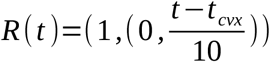

The published auxin-dependent wall mechanics were applied first in all cells^26^. In ccvTIR-positive cells, simulated cvxIAA effect then added a domain-specific modulation to the wall stiffness response, implemented through ccvTIR wall K2 term. The effective cvxIAA term for cell *i* is:

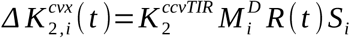

where K is the cvxIAA response strength and the S term is a defined tissue-sensitivity coefficient. The mechanical parameter used by the growth solver was therefore:

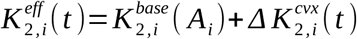

with the baseline term denoting the auxin-dependent response from the previous models. Sensitivity scenarios were encoded with *ccvTissueSensitivity* and compared three radial hypotheses: outer tissues most sensitive, inner tissues most sensitive, or uniform sensitivity across all tissues. Cell division, auxin transport, baseline wall growth, developmental transitions and all non-cvxIAA mechanical parameters were otherwise kept unchanged from the physiological baseline model^26^.

## Author contribution

S.M.D., K.W. and M.F. designed the study. J.E.P. conceptualized the computational modelling. S.M.D., M.S., E.M., P.K. and M.F. performed the experiments. J.E.P. performed the computational modelling. S.M.D. drafted the manuscript, and J.E.P. wrote the computational modelling section. M.F. provided comments on the manuscript. S.M.D., J.E.P. and K.W. edited the manuscript.

## Data Availability

The experimental data generated and analyzed during this study are available in main text and Supplementary Information. The underlying numerical data supporting the figures will be made available as Source Data files upon request from authors. The computational model framework MorphoDynamX used to run models can be found at https://morphographx.org/morphodynamx/.

## Acknowledgement

This work has been funded by Ministerio de Ciencia Innovación y Universidades of Spain (PID2024-155159NB-I00) to K.W., and Severo Ochoa (SO) Program for Centers of Excellence in R&D from the Agencia Estatal de Investigación of Spain [grant CEX2020-000999-S (2022 to 2025) to the CBGP]The authors would like to thank the European Research Council (ERC) for financial support of this work (Grant No. 101125499). S.M.D. was previously supported by the STARS scholarship program of Charles University, Prague, Czech Republic, and is currently supported by a postdoc grant Juan de la Cierva fellowship (Ref. JDC2023-050470-I) funded by the Ministerio de Ciencia, Innovación y Universidades, Spain.

