## Supplementary Information for "Distinct spatial domains of auxin perception control root growth in Arabidopsis"

**Supplementary Figures**

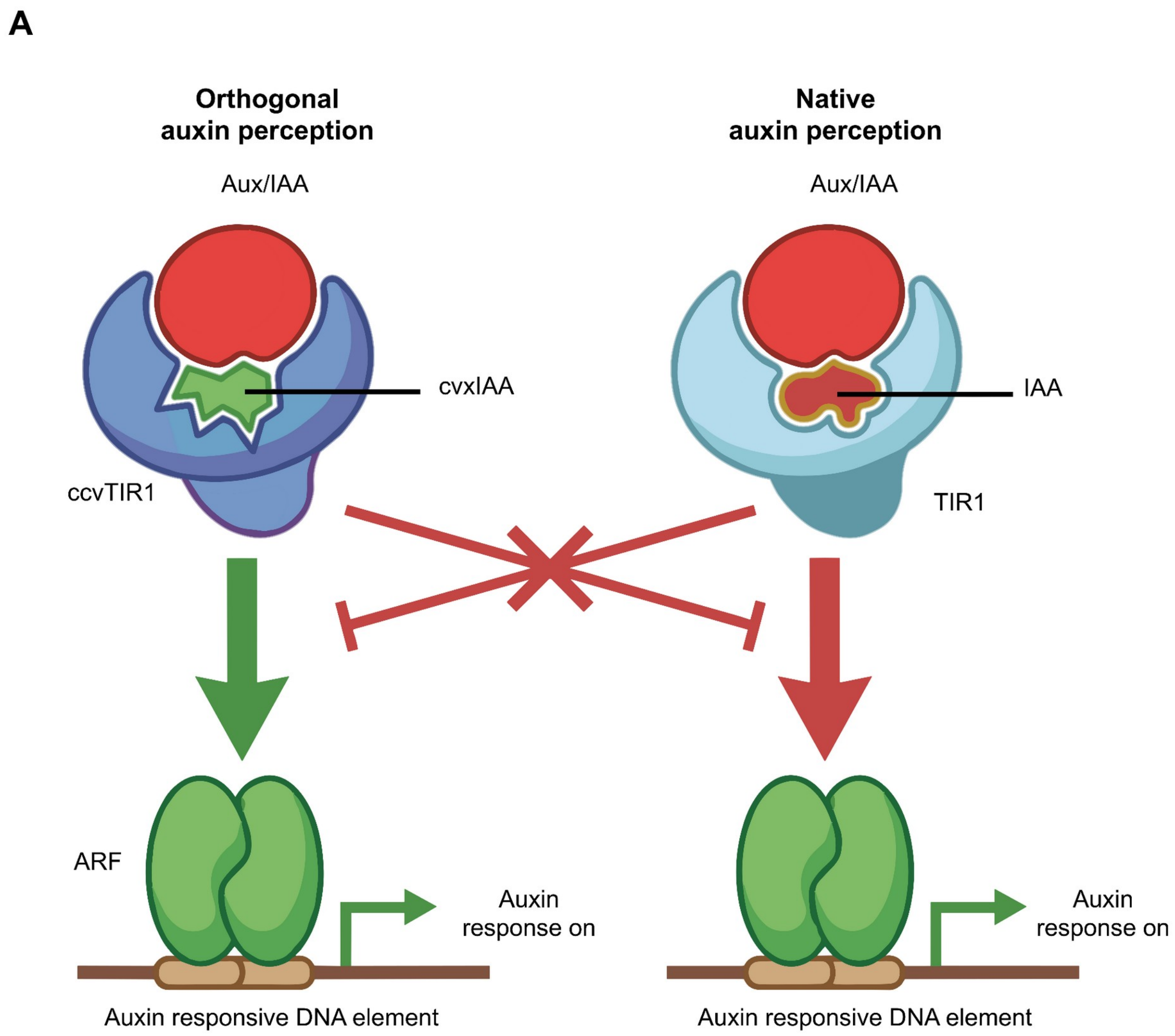


**Supplementary Figure S1. Schematics of orthogonal auxin sensing system.**

**(A)** Schematic of the orthogonal ccvTIR1–cvxIAA pathway (left) and the native TIR1–IAA pathway (right). Each auxin promotes association of its receptor with Aux/IAA. Crossed lines indicate the mismatched receptor and ligand pairs. Aux/IAA degradation releases auxin response factors (ARFs) to activate transcription from auxin-responsive DNA elements.

A

pK890::ccvTIR1-mVenus

B

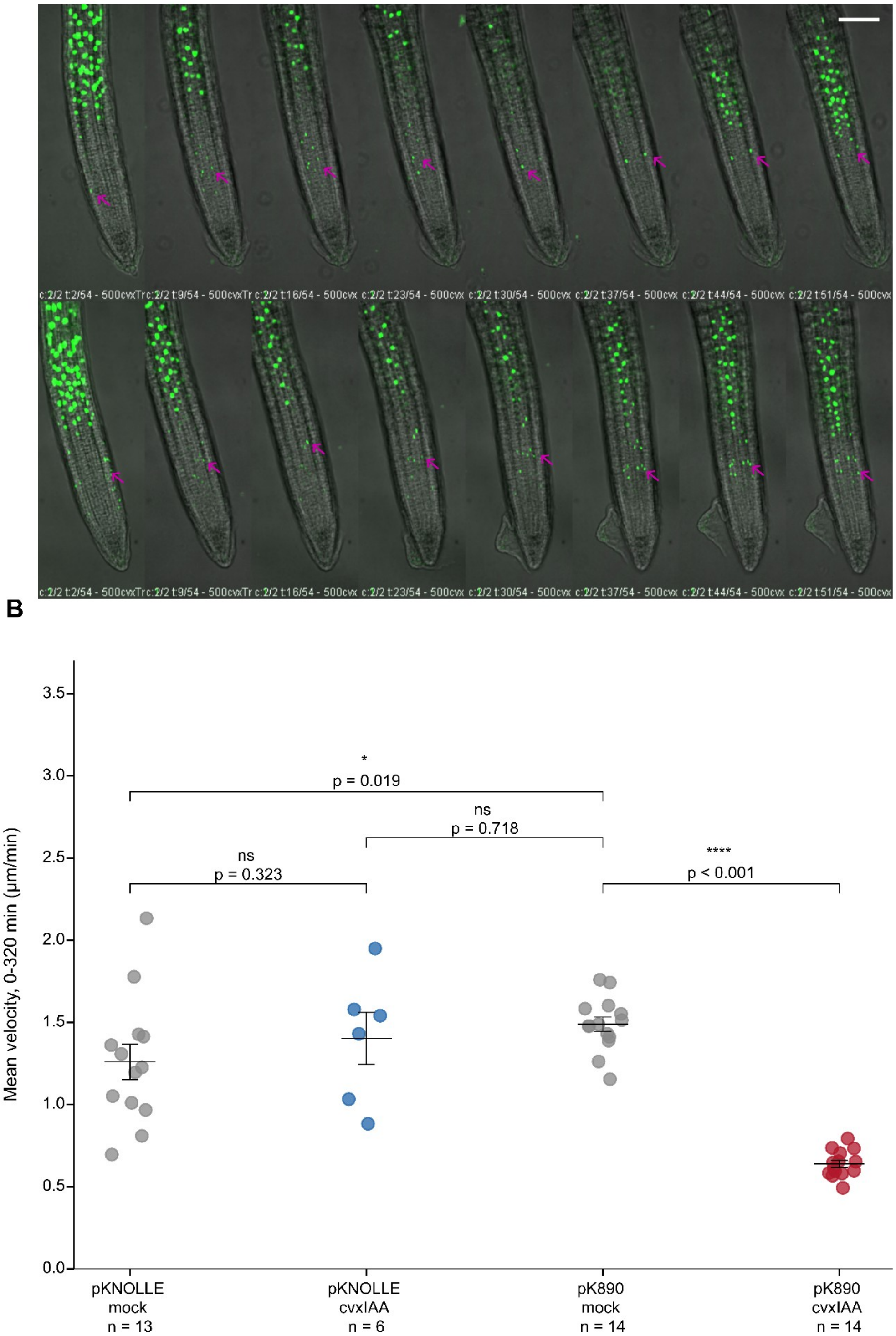

**Supplementary Figure S2. Live imaging of pK890::ccvTIR1–mVenus expression following cvxIAA treatment**

**(A)** Montage showing two pK890::ccvTIR1–mVenus roots treated with 500 nM cvxIAA, demonstrating nuclear expression of pK890::ccvTIR1–mVenus in the elongation zone and distal meristem. The magenta arrows indicate ccvTIR1–mVenus expression in the distal meristem. Images were acquired by live-cell imaging, and the displayed frames were selected after every 2.3 h of imaging. The labels at the bottom of each row indicate the frame number out of a total of 54 frames. Scale bars = 100 μm. **(B)** Per-nucleus mean velocity (averaged over the 0–320 min tracking window; same nuclei as in Fig. 4C, D) compared across all four conditions. Points show individual nuclei; black bars show mean ± SEM. Brackets indicate pairwise Mann–Whitney U tests: mock vs cvxIAA within each genotype (pKNOLLE: $p = 0.32$, ns; pK890: $p < 0.0001$), and cross-genotype comparisons of baseline (pKNOLLE mock vs pK890 mock: $p = 0.019$) and of cvxIAA-treated pKNOLLE cells against untreated pK890 cells ($p = 0.72$, ns). The elongation velocity of cvxIAA-treated pKNOLLE cells is statistically indistinguishable from untreated pK890 cells, despite both pKNOLLE conditions differing from pK890 mock at baseline, indicating that meristem-triggered auxin signaling does not carry over to impair elongation-phase kinetics once cells exit the meristem.

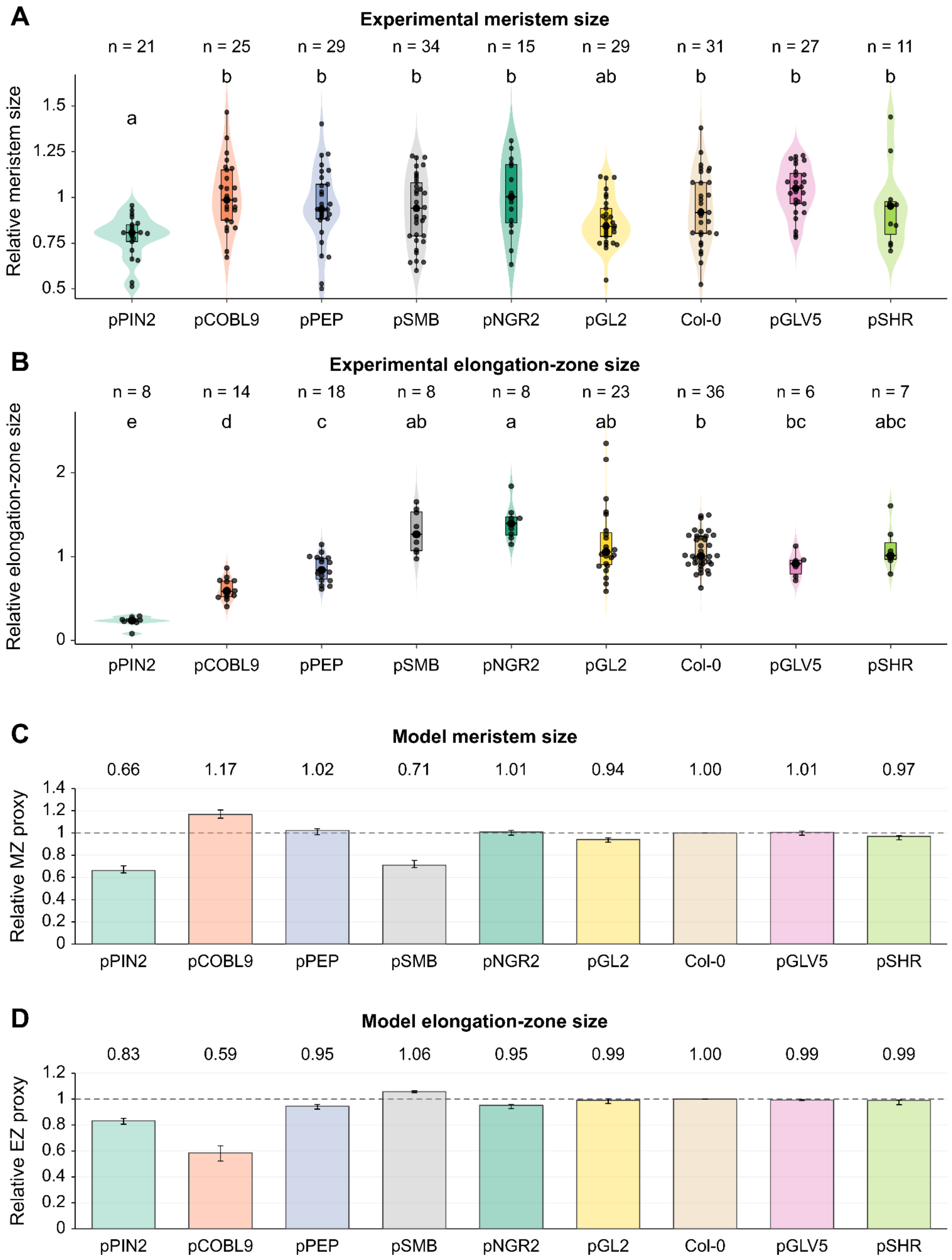
A
Experimental meristem size
n = 21
n = 25
n = 29
n = 34
n = 15
n = 29
n = 31
n = 27
n = 11
a
b
b
b
b
ab
b
b
b
Relative meristem size
1.5
1.25
1
0.75
0.5
pPIN2
pCOBL9
pPEP
pSMB
pNGR2
pGL2
Col-0
pGLV5
pSHR
B
Experimental elongation-zone size
n = 8
n = 14
n = 18
n = 8
n = 8
n = 23
n = 36
n = 6
n = 7
e
d
c
ab
a
ab
b
bc
abc
Relative elongation-zone size
2
1
0
pPIN2
pCOBL9
pPEP
pSMB
pNGR2
pGL2
Col-0
pGLV5
pSHR
C
Model meristem size
0.66
1.17
1.02
0.71
1.01
0.94
1.00
1.01
0.97
Relative MZ proxy
1.4
1.2
1
0.8
0.6
0.4
0.2
0
pPIN2
pCOBL9
pPEP
pSMB
pNGR2
pGL2
Col-0
pGLV5
pSHR
D
Model elongation-zone size
0.83
0.59
0.95
1.06
0.95
0.99
1.00
0.99
0.99
Relative EZ proxy
1.2
1
0.8
0.6
0.4
0.2
0
pPIN2
pCOBL9
pPEP
pSMB
pNGR2
pGL2
Col-0
pGLV5
pSHR

**Supplementary figure S3. Outer tissues exert dominant control over root zonation through auxin signaling.**

**(A)** Quantification of relative meristem size after 20 h treatment with 500 nM cvxIAA in tissue-specific ccvTIR1 lines. Meristem size of each root was normalized to the average mock value of the corresponding genotype. Violin plots represent data distribution, embedded box plots indicate median and interquartile range, and dots represent individual roots. Numbers above each genotype indicate sample size (*n*). Different letters denote statistically significant differences determined by non-parametric ANOVA followed by multiple comparison tests. **(B)** Quantification of relative elongation zone size after 20 h treatment with 500 nM cvxIAA. Elongation zone size was normalized to the average mock value of the corresponding genotype. Violin plots show the distribution of measurements, embedded box plots indicate median and interquartile range, and dots represent individual roots. Numbers above each genotype indicate sample size (*n*). **(C, D)** Model proxies for meristem size (C) and elongation-zone length (D) in simulated roots. Meristem size was estimated from the distance between the lateral root cap boundary and the quiescent center. Elongation-zone length was estimated from the span of EZ epidermal cells along the root axis. Each measurement was divided by the corresponding Col-0 value at the same time point. Bars show the mean ratio over model time, and whiskers span the lowest and highest ratios.

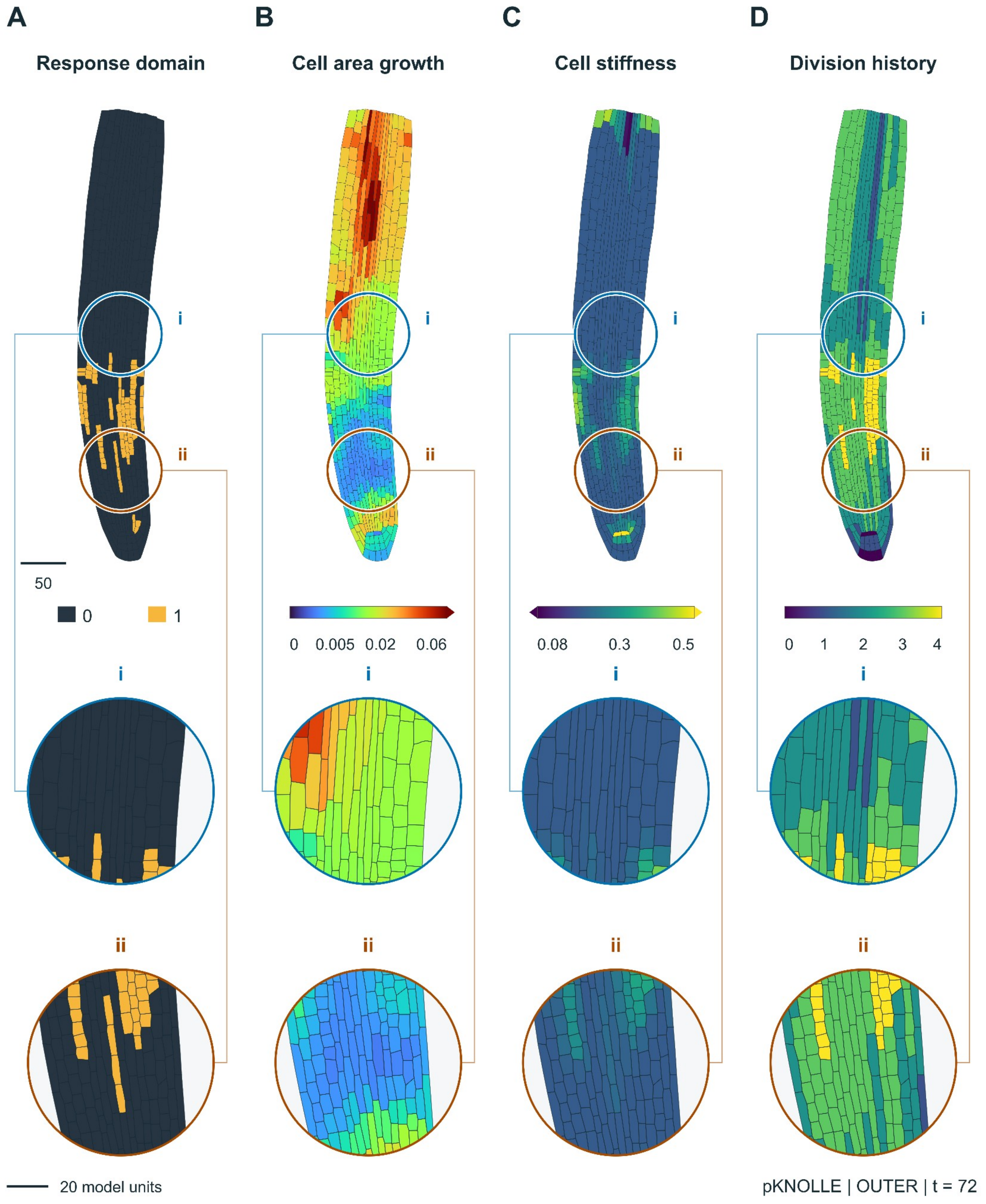


**Supplementary Figure S4. Quantitative metrics to evaluate cell growth and mechanics in the model** Each lens highlights a different feature of the same simulated pKNOLLE root. All four views show the root at model time 72, under a condition that assigns greater auxin sensitivity to the outer tissues. Insets *i* (blue)

and *ii* (brown) show matched regions across all four panels at approximately 2.3× magnification. Their centers are (0, 250) and (0, 100), respectively, with a radius of 45 model length units. Coordinates are relative to the root's horizontal center and lowest visible point. Insets preserve cell identities, triangulation, geometry, and field values. Scale bars, 50 model length units (whole root) and 20 model length units (insets). **(A)** Binary ccvTIR1 response mask. Yellow marks cells where the model applies the ccvTIR response (1), and dark blue marks cells outside the domain (0). **(B)** Cell-level map of growth in the simulated root. Colors show how quickly each cell's area increases relative to its size. Dark blue indicates little or no growth, and red indicates rapid expansion. **(C)** Spatial distribution of cell stiffness along the root axis and across tissue layers. Dark purple indicates lower stiffness, and yellow indicates higher stiffness. Higher values indicate greater resistance to deformation as cells grow. **(D)** Cell-level map of division history. The color of each cell indicates the number of divisions in its lineage. Dark purple corresponds to zero divisions and yellow to four on the displayed scale.

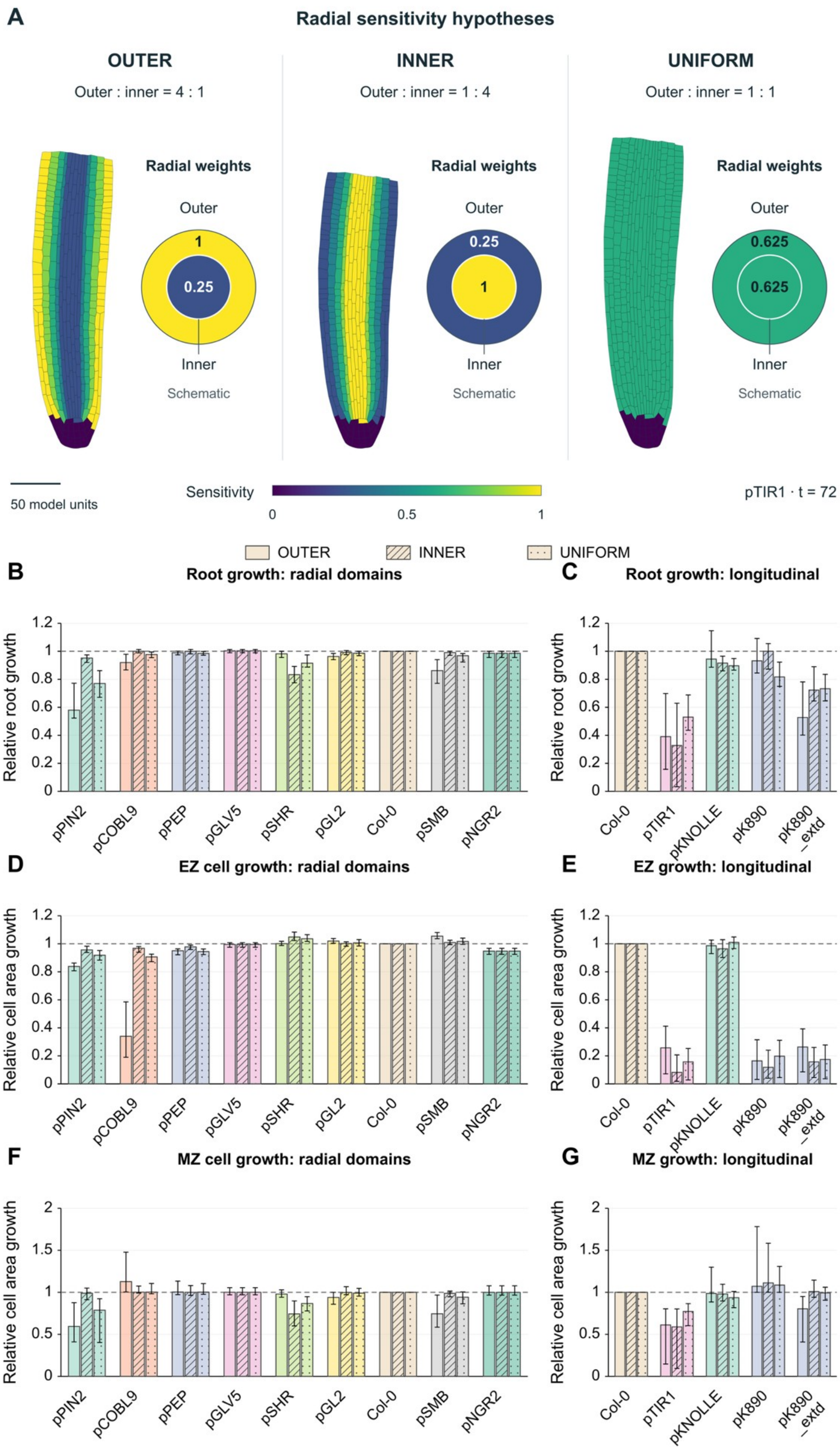

A
Radial sensitivity hypotheses
OUTER
Outer : inner = 4 : 1
INNER
Outer : inner = 1 : 4
UNIFORM
Outer : inner = 1 : 1
Radial weights
Outer
Inner
Schematic
1
0.25
0.625
50 model units
Sensitivity
0
0.5
1
pTIR1 · t = 72
OUTER
INNER
UNIFORM
B
Root growth: radial domains
C
Root growth: longitudinal
Relative root growth
D
EZ cell growth: radial domains
E
EZ growth: longitudinal
F
MZ cell growth: radial domains
G
MZ growth: longitudinal
Relative cell area growth
pPIN2
pCOBL9
pPEP
pGLV5
pSHR
pGL2
Col-0
pSMB
pNGR2
pTIR1
pKNOLLE
pK890
pK890 _extd

**Supplementary Figure S5. Radial auxin sensitivity and simulated growth responses.**

**(A)** ccvTIR sensitivity maps and radial weight schematics at step 2400 (model time 72). Outer and inner weights are 1 and 0.25 for outer, 0.25 and 1 for inner, and 0.625 and 0.625 for uniform. The rings and discs represent tissue identities and assigned weights. The maps share a spatial scale and have aligned tips. The maps and schematics use the same 0–1 color scale. Cells outside the receptor domain retain zero cvxIAA sensitivity. Scale bar, 50 model length units. **(B, C)** Relative whole-root growth rates for simulations in which auxin perception is restricted to different root tissues The three bars for each response domain compare the sensitivity schemes while keeping the responding tissues fixed. **(D, E)** Relative rates of cell area expansion within the elongation zone (EZ). Panel D compares the tissue-specific response domains in B, and panel E compares the longitudinal response domains in C. **(F, G)** Relative rates of cell area expansion within the meristematic zone (MZ) for the same tissue-specific (F) and longitudinal (G) response domains. Panels D–G quantify the expansion of cells within each zone. Solid, diagonally hatched, and dotted bars represent outer, inner, and uniform sensitivity scenarios, respectively. At each time point, growth rates were divided by the corresponding Col-0 rate under the same sensitivity scheme. The dashed line marks the Col-0 reference (1). Bars show the median ratio over model time 15–72, and whiskers span the 25th to 75th percentiles across time.

**Supplementary Movie Legends**

**Supplementary Movie S1. Longitudinal domains of auxin-dependent growth regulation.**

Col-0, pTIR1, pKNOLLE, pK890, and pK890_extd are shown side by side. Each section follows the same simulations from model time 0.03 to 72. **(A)** ccvTIR1 response domains in the simulated roots. Yellow marks cells with an active response, and dark blue marks inactive cells. In pK890_extd, the response domain extends into the lateral root cap. **(B)** Spatial distribution of cell area growth. Colors indicate how quickly each cell's area increases relative to its size. Values below the roots give the whole-root growth rate relative to Col-0 at the same model time. **(C)** Spatial distribution of cell stiffness along the root axis and across tissue layers. Higher values indicate greater resistance to deformation. **(D)** Simulated root zonation. Blue marks the meristematic zone, and pink marks the elongation zone. Values below the roots show the distance from the LRC boundary to the QC, normalized to Col-0. Scale bar, 50 model length units.

**Supplementary Movie S2. Long-term growth dynamics of the pK890 line under mock and cvxIAA treatment.**

Seedlings of the pK890 line were grown under mock conditions or treated with 500 nM cvxIAA in a customised growth chamber mounted on a vertical-stage spinning-disk microscope. Following cvxIAA treatment, pK890 fluorescence appeared closer to the root tip compared with the mock condition, indicating a reduction in the extent of the meristematic domain. In addition to its expression in elongating cells, localised pK890 fluorescence was also visible in the lateral root cap (LRC). Root growth and cell elongation were visibly reduced following cvxIAA treatment, consistent with the reduced cell velocity observed in the nuclear tracking analysis (Fig. 4C).

**Supplementary Movie S3. Long-term growth dynamics of the pKNOLLE line under mock and cvxIAA treatment.**

Seedlings of the pKNOLLE line were grown under mock conditions or treated with 500 nM cvxIAA in a customised growth chamber mounted on a vertical-stage spinning-disk microscope. Following cvxIAA treatment, the reduction in the meristematic domain was accompanied by a decrease in mVenus fluorescence, consistent with a reduction in the pKNOLLE expression domain. In contrast to the pK890 line, cell elongation appeared largely unaffected during the imaging period. This observation is consistent with nuclear tracking analysis showing comparable cell velocities under mock and cvxIAA conditions (Fig. 4D).

**Supplementary Movie S4. Tissue-specific auxin responses across radial domains.**

The nine simulations follow the order in Fig. 5C: pPIN2, pCOBL9, pPEP, pGLV5, pSHR, pGL2, Col-0, pSMB, and pNGR2. All four sections replay the same simulation times at a common spatial scale.

**(A)** ccvTIR1 response domains in the simulated roots. Yellow marks cells with an active response, and dark blue marks inactive cells. **(B)** Spatial distribution of cell area growth. Colors indicate how quickly each cell's area increases relative to its size. Values below the roots give the whole-root growth rate relative to Col-0 at the same model time. **(C)** Spatial distribution of cell stiffness along the root axis and across tissue layers. Higher values indicate greater resistance to deformation. **(D)** Simulated root zonation. Blue marks the meristematic zone, and pink marks the elongation zone. Values below the roots give the meristem-size estimate relative to Col-0 at the same model time. Scale bar, 50 model length units.

**Supplementary Movie S5. Effects of outer and inner tissue sensitivity on root growth.**

Side by side comparisons of inner, outer and uniform sensitivity schemes are shown for Col-0, pPIN2, pK890_extd, and pTIR1, the conditions showed the largest differences among schemes in median whole-root growth rate relative to Col-0. The opening frames show the assigned auxin sensitivities at model time

15 on a common color scale from 0 to 1. Outer assigns relative sensitivities of 1 to outer tissues and 0.25 to inner tissues. Inner reverses these values, and Uniform assigns 0.625 to both tissue groups. The time lapse shows local cell area growth through model time 72. Colors indicate how quickly each cell's area increases relative to its size. Values below the roots give the whole-root growth rate relative to Col-0 under the same sensitivity scheme at the displayed time. Scale bar, 50 model length units.
